# Biocompatible designated Resin-3D-printed polymers exhibit reproductive toxicity prevented by Parylene-C

**DOI:** 10.64898/2026.06.05.730268

**Authors:** Hannes Campo, Uyen Tran, Yiru Zhu, Hoi Chang Lee, Francesca E. Duncan

**Affiliations:** Department of Obstetrics and Gynecology, Feinberg School of Medicine, Northwestern University, Chicago, IL, 60611; Buck Institute for Research on Aging, Novato, CA, 94945

**Keywords:** 3D printing, biocompatibility, oocyte, reprotoxicity, parylene-c, microphysiological systems, resin, new approach methodologies

## Abstract

Resin three-dimensional (3D) printing is an increasingly popular manufacturing and prototyping method used to create microphysiological systems (MPS), but resin cytotoxicity significantly hinders its adoption, especially when sensitive cell models are incorporated. The mammalian oocyte and early preimplantation embryo consist of cells that are highly sensitive to toxicants and thus represent stringent cell-based models for biocompatibility testing. We developed a Multi-Endpoint Oocyte Safety Assay (MEIOSA) to evaluate the biocompatibility of four ISO 10993 biocompatible BioMed resins (Clear, Durable, Elastic 50A, and Flex 80A). MEIOSA assesses the viability, morphology, meiotic stage, and meiotic spindle morphology of the oocyte after *in vitro* maturation (IVM). Oocytes were *in vitro* matured in plate inserts 3D printed with the four BioMed resins. Oocytes cultured in rigid resins (Clear and Durable) or elastomeric resins (Elastic 50A, and Flex 80A) exhibited impaired meiotic progression and complete oocyte degeneration, respectively, relative to controls cultured in polystyrene which matured normally. To determine whether such cytotoxicity could be prevented, we coated the resin inserts with a 5 µm impermeable Parylene-C (PC) barrier. PC coating completely rescued the degeneration and meiotic maturation defect phenotypes for all resins. Remarkably, when the most cytotoxic material (Flex 80A) was coated with PC, the resulting eggs were fertilization-competent and produced embryos capable of normal preimplantation development via *in vitro* fertilization. Our findings demonstrate that standardized viability-based biocompatibility tests do not identify cytotoxic effects for all cell types and establish MEIOSA as a high sensitivity test model to robustly evaluate biomaterial biocompatibility. Furthermore, PC coating prevents the toxic effects of all resin-3D-printed materials tested, opening up a new toolbox to create MPS compatible with reproductive, and by extension, other sensitive cell cultures.

**Graphical abstract:** 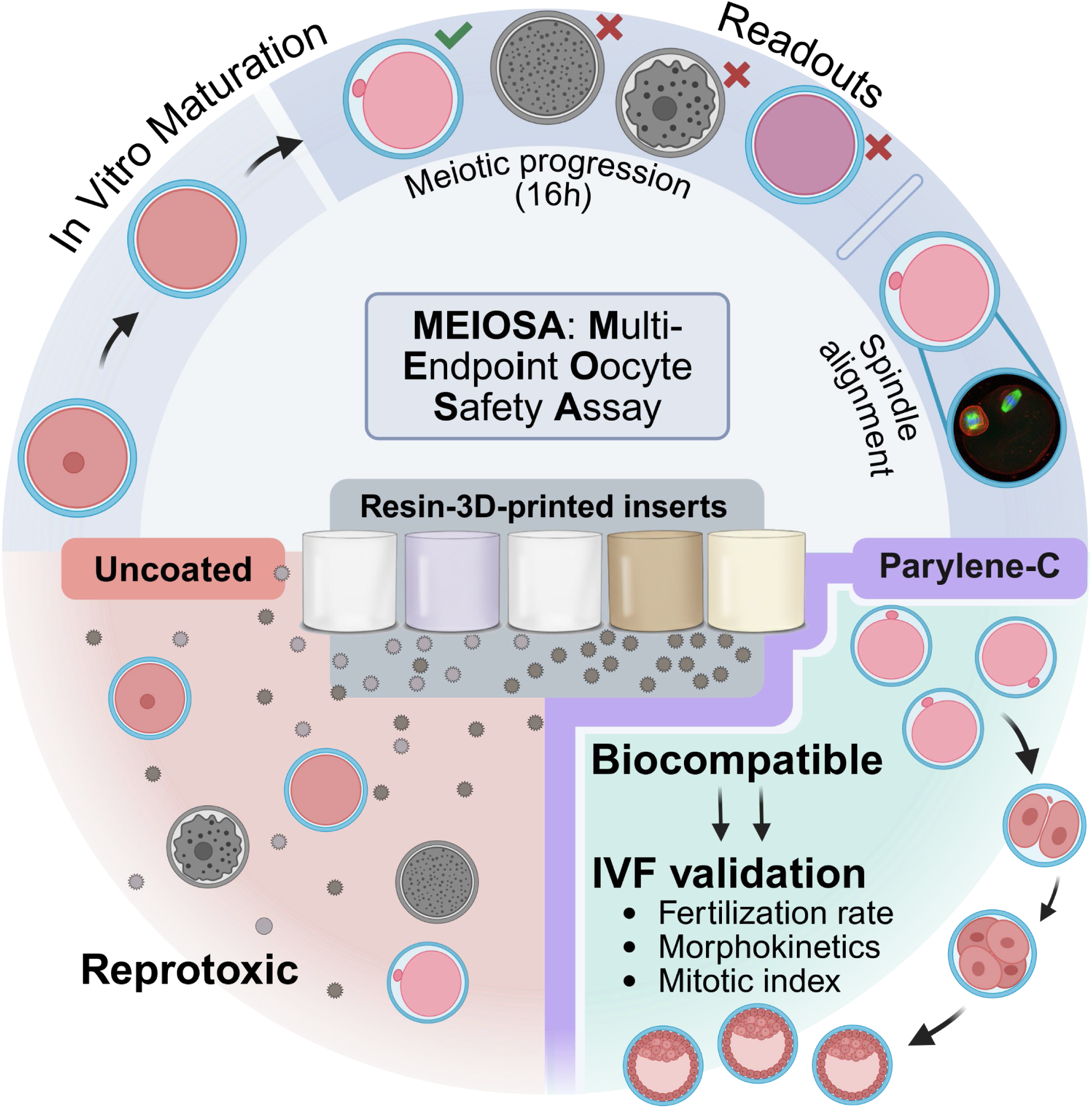

## 1. Introduction

Accurately modeling human physiology *in vitro* is essential for biomedical research, toxicology studies, and drug development.^[1–3]^ Traditional static two-dimensional cultures are unable to recapitulate cellular complexity, biochemical cues, and tissue interactions, which hinders the translation of insights from basic research into mechanisms of health and disease, and finally to clinical applications.^[4]^ To address these limitations, microphysiological systems (MPS) were engineered as *in vitro* models that incorporate physiologically relevant microenvironments of cells and tissues. MPS are a central part of the era of human-relevant new approach methodologies (NAMs), which based on the FDA Modernization Act 2.0/3.0, may be used to reduce or replace conventional animal studies where appropriate.^[5–8]^ Hence, it is expected that the adoption and refinement of these technologies will increase as our reliance on animal and 2D models decreases, and basic research and clinical applications move towards human and personalized MPS.^[3,4,9]^

Key considerations during the design and implementation of MPS are material selection, fabrication, and biocompatibility.^[10]^ Vat-polymerization (VP), also known as resin three-dimensional printing, is an additive manufacturing technique where sequential layers of photopolymerizable resins are light-cured. VP is used in dental applications,^[11]^ medical devices,^[12]^ and MPS.^[13–17]^ Sequential light curing can be performed with high precision, and the most common methods utilize lasers (stereolithography apparatus, SLA) or light projection (digital light processing, DLP).^[16–19]^ VP offers several benefits compared to other manufacturing technologies, including affordability, precise control over complexity and design parameters, rapid iteration, and scalable manufacturing. Furthermore, multiple materials and hydrogels with varying mechanical and biochemical properties can be used.^[16–19]^

However, a critical limitation of using resin-3D-printed materials for culture applications is the release of cytotoxic leachates, such as monomers, surfactants, or photochemical compounds.^[11,20–22]^ The toxicity of these leachates is commonly determined by using somatic cells with viability-based endpoints, which also forms the basis for standard *in vitro* testing during ISO 10993 validation. The ISO 10993 suite of standards specifies the requirements and test methods for evaluating the safety of medical device materials but does not take into account whether a material can be used for MPS or highly sensitive cultures.^[23]^ In fact, multiple studies demonstrate that leachates of resins deemed to be “biocompatible” induce cell death, cell detachment, sub-cytotoxic perturbations, and endocrine effects, even after aggressive post-processing protocols.^[11,24–26]^ Furthermore, there is no general consensus as to which biocompatibility standards are relevant for resin-3D-printed MPS.

The goals of this study were threefold: to (1) develop a multi-endpoint oocyte safety assay (MEIOSA) for standardized and rapid biocompatibility testing in a highly sensitive cell model compatible with MPS applications, (2) use MEIOSA to evaluate a line of commercially available ISO 10993 validated biocompatible resins with a wide range of mechanical properties (Clear, Durable, Elastic 50A, and Flex 80A), and (3) test if Parylene-C (PC) coating, a common and affordable FDA-approved USP Class VI polymer able to form micron-thin impermeable barriers, improves biocompatibility for all 3D-printed-resins.

The MEIOSA assay leverages the highly sensitive maturation process of the mammalian oocyte that occurs at the time of ovulation. During ovulation, prophase I-arrested oocytes complete meiosis I and arrest in metaphase II (MII), which is a high-risk event for chromosomal reorganization. This biological pathway is highly sensitive to material-specific leachates and other environmental contaminants, such as the endocrine disruptor BPA and its variants, leading to aneuploidy.^[11,27–29]^ Furthermore, oocyte maturation can be induced *in vitro*, and key maturation events can be monitored morphologically, which provides quantifiable readouts after 16-hours of culture. Oocyte quality can then be assessed further using immunocytochemistry, as chromosomes also have to accurately align on the metaphase plate within the meiotic spindle.^[30,31]^ Hence, oocyte morphology, spindle formation and chromosome alignment give highly stringent single-cell level readouts for *in vitro* cytotoxicity assessment.^[11]^ Using MEIOSA, we evaluated a new line of ISO 10993 biocompatible (BioMed) resins (Clear, Durable, Elastic 50A, and Flex 80A) and one control resin (Clear v4). Maturation in all biomaterials resulted in disrupted meiotic maturation or induced cell death. However, PC coating completely prevented toxicity for all resins. The resulting eggs following IVM in the most toxic biomaterial coated with PC were able to be fertilized and undergo normal preimplantation embryo development following *in vitro* fertilization (IVF). Our findings demonstrate that viability-based testing fails to comprehensively evaluate the *in vitro* biocompatibility of 3D printed resins which poses significant safety and toxicity risks especially for highly sensitive cells such as the germline which can give rise to the next generation. The MEIOSA assay, based on the mammalian oocyte, is thus a critical model system for evaluating biocompatibility using the most sensitive cell types as the highest safety barometer. We also show that PC coating is an accessible way to prevent reprotoxicity, broadening the number of biomaterials that can be used in a biocompatible manner to build MPS.

### 2. Results

#### 2.1. The Multi-Endpoint Oocyte Safety Assay (MEIOSA)

##### 2.1.1. MEIOSA enables assessment of BioMed material biocompatibility using multiple oocyte endpoints

To evaluate a new line of biocompatible resins (BioMed, BM), we designed inserts that fit into 24-well plates and allowed for oocytes to be seeded, optical imaging to occur, and leachates to diffuse into the media (**Figure 1A-B**). In total five different resins were selected to include a wide spectrum of mechanical properties and biocompatibility claims, and these were 3D printed and post-processed according to manufacturers protocols (**Figure 1B-C**). We used four BM resins (BM Clear, BM Durable, BM Elastic 50A, and BM Flex 80A) which adhere to ISO 10993 biocompatibility standards of medical devices for human use. Of note, BM Clear passed ISO-10993-3 which encompasses tests for genotoxicity, carcinogenicity, and reproductive toxicity (**Figure 1C**). BM Clear and BM Durable are rigid materials, intended for long-term skin or mucosal membrane contact, whereas BM Elastic 50A and BM Flex 80A are elastomeric, used for flexible biomedical device components. Clear V4, which has no biocompatibility certification and verified cytotoxicity,^[32]^ was used as a control for the assay (**Figure 1C**).

**Figure 1.**
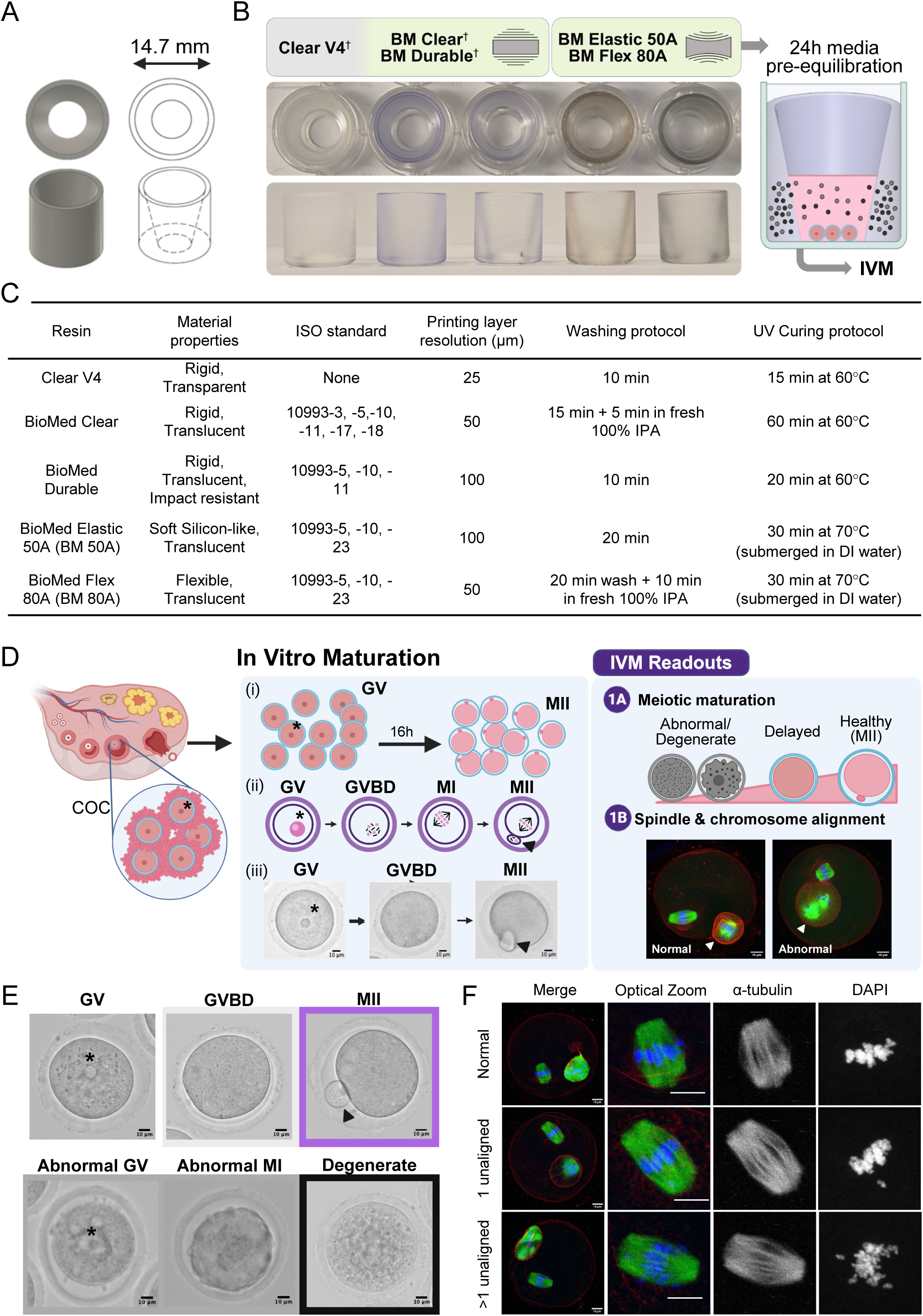
Determining the biocompatibility of 3D-printed resins using the Multi-Endpoint Oocyte Safety Assay (MEIOSA). A) Well inserts fit 24-well culture plates and were designed to allow for easy imaging and co-culture with oocytes. B) In total 5 resins were tested: three rigid (†) and two elastomeric inserts that are non-biocompatible (grey) or adhere to ISO 10993 standards (green). Culture media was pre-equilibrated with inserts for 24 hours prior to oocyte seeding. C) Properties, manufacturing, and post-processing parameters of analyzed resins. ISO standards specify the requirements and test methods for evaluating genotoxicity, carcinogenicity, and reproductive toxicity (ISO 10993-3), *in vitro* cytotoxicity (ISO 10993-5), skin sensitization (ISO 10993-10), identification and quantification of chemical components (ISO 10993-18), systemic toxicity (ISO 10993-11), allowable limits for leachable substances (ISO 10993-17) and potential to produce irritation (ISO 10993-23) of medical device materials. UV: ultraviolet, IPA: isopropyl alcohol, DI; Deionized. D) Oocytes at the germinal vesicle stage (GV, indicated by asterisk) are isolated mechanically from cumulus-oocyte-complexes (COC). (i) During *In vitro* Maturation, GV oocytes progress to the metaphase II (MII) stage over a 16h culture period. (ii) The stages of meiosis moving from GV to MII include the GV breakdown (GVBD), chromosomal reorganization and polar body extrusion (PB, indicated by triangle). (iii) Representative images of oocytes throughout IVM in polystyrene control culture plate demonstrate that meiotic progression can be observed morphologically. After 16 hours, IVM readouts such as meiotic maturation, where the oocyte is abnormal, degenerate, delayed or healthy, can be determined (1a), followed by spindle chromosome alignment (1b). E) Representative images for classifications of oocyte meiotic maturation phenotypes (germinal vesicle stage (GV*), germinal vesicle breakdown/metaphase I (GVBD), metaphase II (MII), abnormal (at MI or MII stage), or degenerate. F) Representative confocal microscopy images of normal and abnormal chromosome and spindle architecture. Actin (rhodamine phalloidin, red), α-tubulin (green), and DNA (DAPI, blue and greyscale insert). PB indicated by triangle.

To test whether the resins were biocompatible with mammalian oocytes, we established the MEIOSA assay to monitor meiotic progression of oocytes during IVM. Fully grown oocytes arrested in prophase of meiosis I were isolated and denuded from cumulus oocyte complexes (COCs) collected from antral follicles from hyperstimulated mice (**Figure 1D**). Prior to seeding oocytes, we incubated media in the inserts for a 24 h pre-equilibration leaching period (**Figure 1B**). Oocytes were then *in vitro* matured for 16 h during which time they progressed from prophase of meiosis I to metaphase of meiosis II (**Figure 1D**). These transitions through meiosis can be visualized morphologically by transmitted light microscopy (**Figure 1D-E**). Prophase I arrested cells are characterized by an intact nucleus or germinal vesicle (GV). Cells that have progressed to pro-Metaphase I or Metaphase I have undergone germinal vesicle breakdown (GVBD) and lack a GV but have not yet extruded a polar body. Cells that have fully matured and reached Metaphase of Meiosis II (MII) are characterized by the extrusion of a polar body (**Figure 1E**). At the end of culture, healthy versus abnormal or degenerate oocytes were scored based on cytoplasmic morphology (**Figure 1D-E**). Oocytes with abnormal morphologies are characterized by cellular aggregates, vacuoles, and membrane protrusions, whereas degenerate oocytes have a flattened and transparent appearance (**Figure 1E**). For all healthy oocytes, meiotic stage was scored. Normal MII eggs are characterized by a bipolar spindle and chromosomes tightly aligned on the metaphase plate, and this process is highly sensitive to perturbation.^[29]^ Therefore, we also assessed meiotic spindle structure in all resulting MII eggs (**Figure 1D**). Alignment of chromosomes on the metaphase plate for oocytes that reached MII was categorized as normal, with one misaligned chromosome, or with multiple misaligned chromosomes (**Figure 1F**). Importantly, we have previously used and validated a similar assay to determine the impact of leachates on oocytes from dental resins.^[11]^

##### 2.1.2. Resins deemed biocompatible are cytotoxic to oocytes with elastomeric materials exhibiting more severe effects compared to rigid resins

To determine the impact of the BioMed resins on oocyte meiotic progression, we *in vitro* matured oocytes in inserts made of the various resins using the MEIOSA pipeline (**Figure 2**). Despite IVM occurring only over the course of hours, clear distinctions were apparent between the biomaterials. The majority of oocytes matured in control polystyrene wells reached MII as expected (86.7 ± 9.4%). No oocytes matured in Clear V4 reached MII, with most exhibiting abnormal phenotypes (98.3 ± 2.4%), presenting abnormal membrane morphology and arrest in GVBD (**Figure 2A-B**). Similar findings were observed with the rigid BM materials, with no cells reaching the MII stage and instead abnormal phenotypes, including cellular aggregates, observed in 85.0 ± 11.8% and 93.3 ± 4.7% of oocytes cultured in BM Clear and BM Durable, respectively (**Figure 2A-B**). These findings are consistent with previous observations of resin-induced meiotic disruption.^[11]^ BM Elastic 50A (BM 50A) and BM Flex 80A (BM 80A) exhibited a more cytotoxic profile compared to the rigid inserts, with 100% cellular degeneration observed with both materials (**Figure 2A-B**). Notably, many degenerate oocytes retained GV-like structures, indicating that cell death occurred prior to or early during meiotic resumption and, therefore, rapidly after exposure. Immunofluorescence analysis confirmed normal bipolar spindle morphology with chromosomes aligned on the metaphase plate in control oocytes matured under standard conditions (**Figure 2C**). However, chromosome organization of oocytes cultured with rigid inserts (Clear V4, BM Clear and BM Durable) more closely resembled Prophase I and GVBD configurations (**Figure 2C**). Spindle and chromosome morphology were not assessed for elastomeric conditions due to complete degeneration.

**Figure 2.**
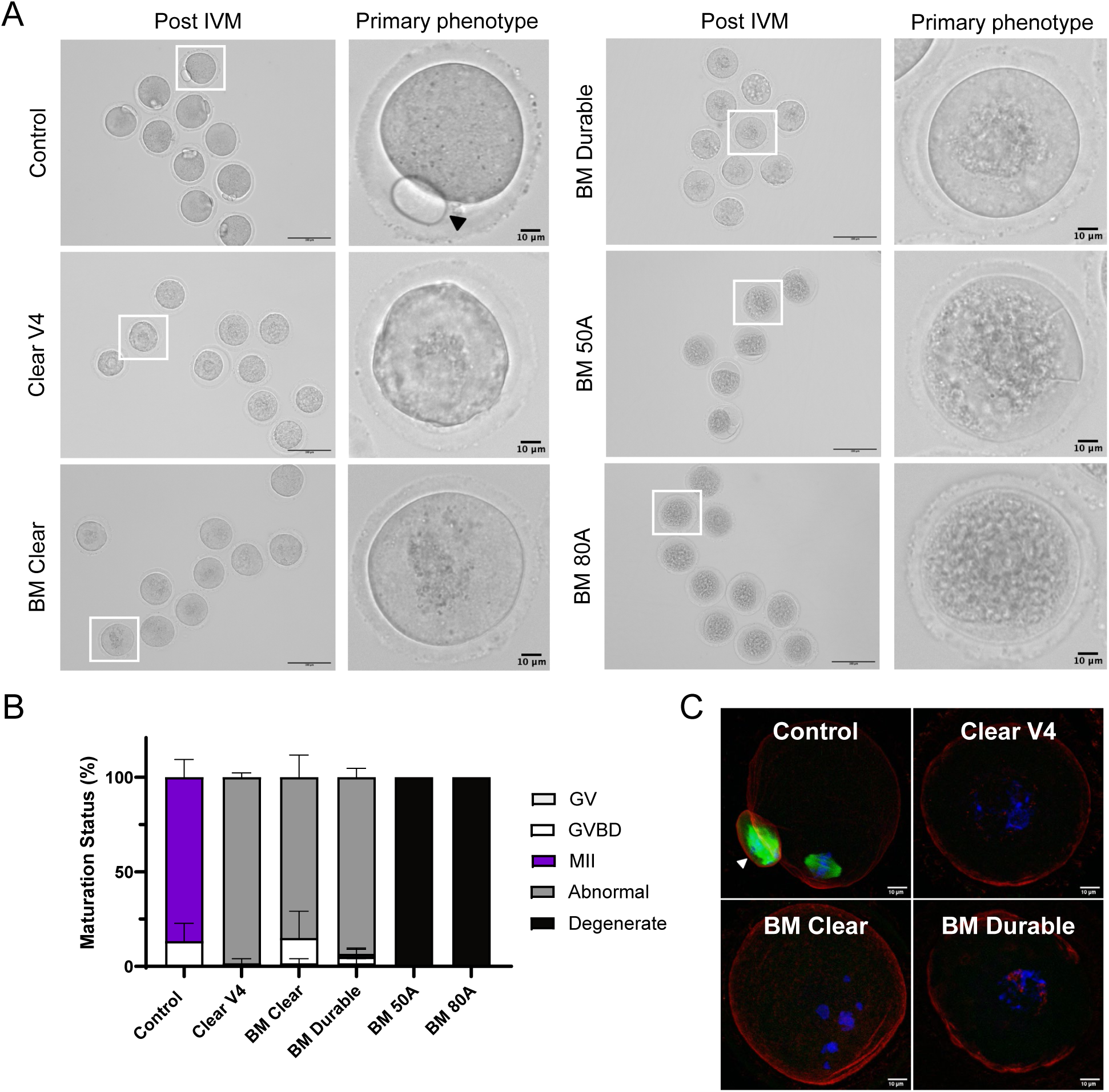
IVM readouts demonstrate a range of cytotoxicity induced by uncoated 3D-printed culture inserts. A) Representative images of oocytes cultured with uncoated inserts after IVM (left, scale bar: 100 µm) with primary phenotypes (right, scale bar: 10 µm). B) Resulting meiotic progression incidence for all materials compared to control. C) Representative z-projections of chromosome and spindle architecture found for control and abnormal phenotypes. Actin (rhodamine phalloidin, red), α-tubulin (green), and DNA (DAPI, blue). PB indicated by triangle, scale bar: 10 µm. For each culture condition a total of 60 oocytes was used (10 oocytes per culture well, three technical and two biological repeats).

##### 2.1.3. Parylene-C coating of rigid materials rescues abnormal phenotypes

Though BM Clear, in some instances, can pass MEIOSA biocompatibility standards, OCP efficiency can vary and may not be useful if other material properties are required in the MPS design. Parylene-C (PC) is a well-suited solution for this purpose, as it can form micron-thin and impermeable barriers, blocking leachates to enter the culture medium.^[33,34]^ Hence, we investigated if a 5 µm thick conformal PC coating could improve biocompatibility (**Figure 3A**). As expected for our control in the MEIOSA assay, IVM under standard conditions resulted in an MII incidence of 95.6 ± 1.6%, and assay consistency was validated in Clear V4 with an incidence of abnormal morphology of 95.6 ± 1.6% (**Figure 3B**). Interestingly, the incidence of MII increased from 0% to 73.3% for BM Clear when inserts were used 82 days post-curing compared to 5 and 12 days post-curing, conditions when >90.0% of oocytes exhibited abnormal phenotypes **(Figure S1A-B)**. These findings indicate that the current manufacturer protocol was insufficient to render BM Clear inserts biocompatible and usable for *in vitro* culture of all cell types. This aging-related improvement was not observed for the other rigid materials tested **(Figure S1C)**.

**Figure 3.**
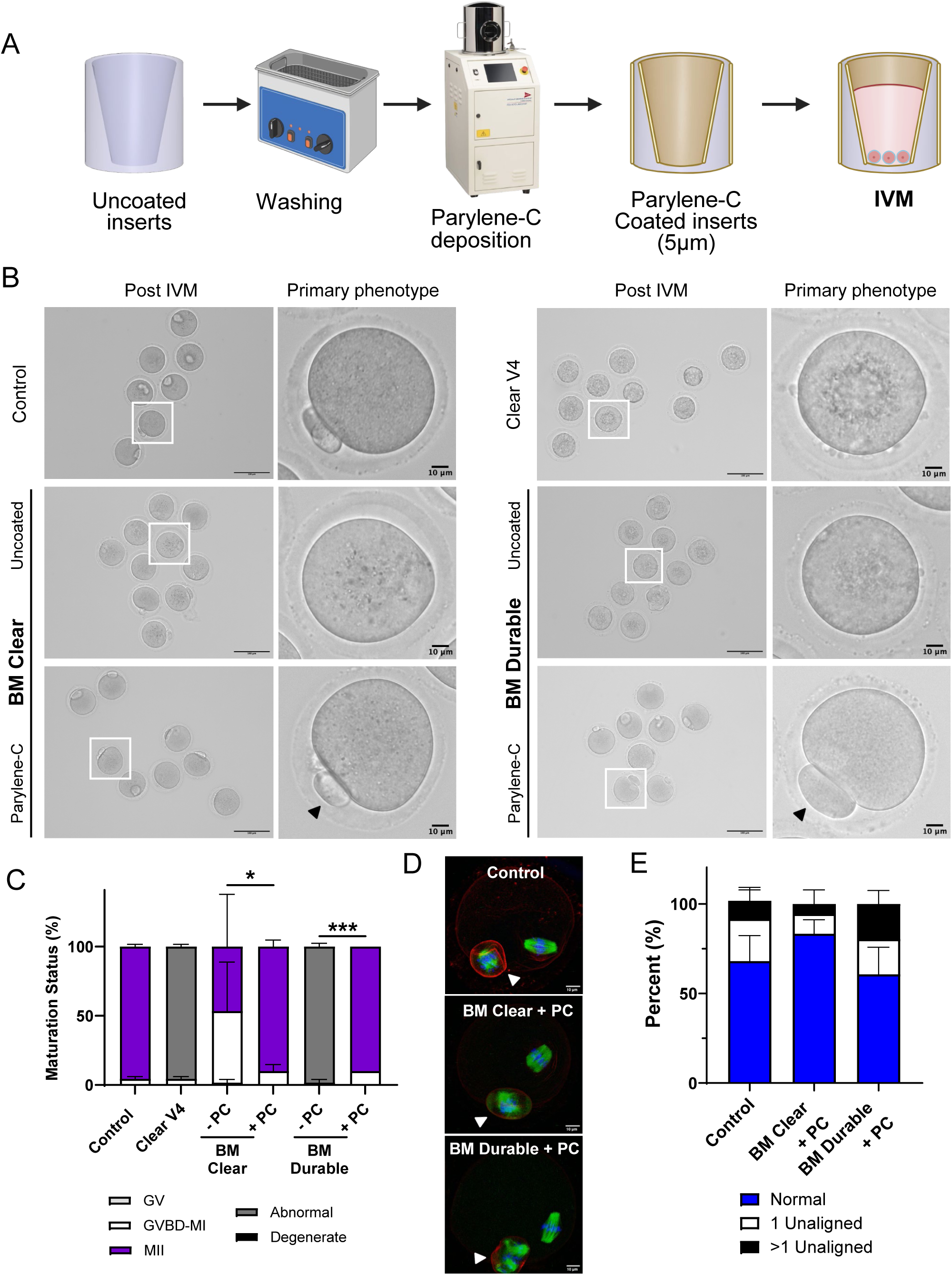
Effect of Parylene-C coating on biocompatibility of rigid BioMed inserts. A) Preparation steps and 5 µm Parylene-C coating procedure prior to oocyte seeding. B) Representative images of primary phenotype observed after IVM for each uncoated and coated material. C) Meiotic maturation incidence. (* and *** represent *P* < 0.05 and *P* < 0.001 when comparing MII rates, respectively). D) Representative z-projections of MII eggs in control and PC coated conditions. Actin (rhodamine phalloidin, red), α-tubulin (green), and DNA (DAPI, blue) were detected by immunocytochemistry. PB indicated by triangle, scale bar: 10 µm. E) Incidence of normal and aberrant chromosome alignment. For each culture condition a total of 60 oocytes was used (10 oocytes per culture well, three technical and two biological repeats).

A common method of improving biocompatibility of 3D printed resins is use of overcuring protocols (OCP) in an effort to remove toxic unreacted monomers.^[35,36]^ To determine whether biocompatibility improvements could be achieved more rapidly, two OCP variations utilizing submerged UV curing (OCP 1) and overnight exposure to heat (OCP 2) were tested. Both approaches resulted in similar incidences of MII relative to the aging OCP protocol used for BM Clear (66.7% for OCP1, and 83.3% for OCP2) **(Figure S1B, D-E)**. The spindle structure and chromosome alignment of eggs matured in all OCP samples for BM Clear were similar to controls matured under standard conditions **(Figure S1F-G)**. These findings demonstrate that MEIOSA readouts can be used to evaluate and rank levels of cytotoxicity of biocompatible rigid and elastomeric resins and that OCP can improve biocompatibility of only BM Clear.

Coating both BM Clear and BM Durable was able to significantly improve the incidence of MII such that the levels were indistinguishable from controls matured under standard conditions (**Figure 3B-C**). For BM Clear, the incidence of MII increased from 46.7 ± 37.7% in the uncoated condition to 90.0 ± 4.7% in the PC coated condition (**Figure 3B-C**). The increased standard deviation observed in the uncoated condition was attributable to aging effects in BM Clear (**Figure S1A-B**). The effects of PC coating on the biocompatibility of BM Durable were more striking, with MII rates improving from 0.0 ± 0% to 90.0 ± 0% (**Figure 3C**). Chromosome alignment analysis confirmed the effectiveness of PC further, with 83.3 ± 7.9% for BM Clear + PC and 60.7 ± 15.2% for BM Durable + PC presenting normal alignment, which was not significantly different from control (68.2 ± 14.2%) (**Figure 3D–E**). These results demonstrate that PC coating renders rigid 3D-printed resins biocompatible with oocyte maturation.

##### 2.1.4. Parylene-C coating of elastomeric materials rescues oocyte degeneration

We next investigated if PC coating could also improve the biocompatibility of elastomeric materials, which were the most cytotoxic (**Figure 4A**). Using the same methodology, PC coating was able to achieve complete reversal of the toxic effects of BM 50A and BM 80A on oocyte survival and development (**Figure 4A-B**). In both conditions, the primary oocyte phenotype following IVM shifted from 100% degenerate to > 90% MII (Figure 4B). The MII incidence for controls matured under standard conditions was 95.6 ± 1.6% compared to 93.3 ± 9.4% for BM 50A + PC and 96.7 ± 4.7% for BM 80A + PC. Furthermore, chromosome alignment analysis showed a normal alignment for the majority of oocytes in BM 50A + PC and BM 80A + PC conditions, at 50.6 ± 18.5% and 64.6 ± 6.5%, respectively. Once again, there was no significant difference between PC-coated materials and control which was at 68.2 ± 14.2% (**Figure 4C-D**). These results demonstrate that PC coating blocks toxic leachates over the combined 24 h pre-equilibration and 16 h IVM periods, completely preventing oocyte cytotoxicity of elastomeric inserts. Of note, the type of resin and the subsequent PC coating procedure had measurable effects on the width of the insert, and these discrepancies should be taken into account when incorporating PC coated 3D printed components in MPS designs **(Figure S2A-B)**.

**Figure 4.**
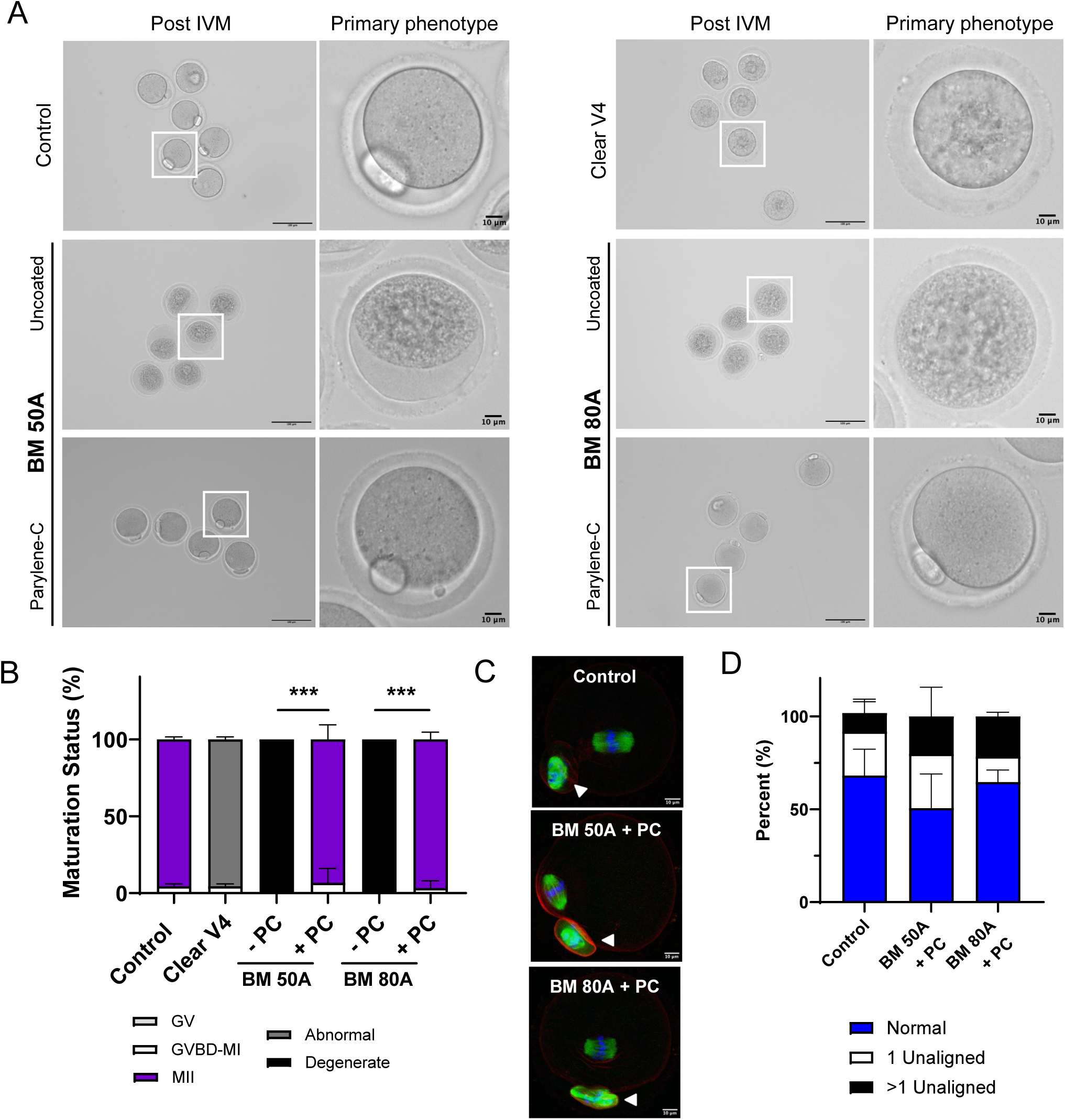
Effects of Parylene-C coating on biocompatibility of cytotoxic elastomeric BioMed inserts. A) Representative images of primary phenotype observed after IVM for each uncoated and coated material. B) Meiotic maturation incidence. *** represents *P* < 0.001 when comparing MII rates. C) Representative z-projections of phenotypes of MII eggs in control and PC coated conditions. Actin (rhodamine phalloidin, red), α-tubulin (green), and DNA (DAPI, blue) were detected by immunocytochemistry. PB indicated by triangle, scale bar: 10 µm. D) Incidence of normal and aberrant chromosome alignment. For each culture condition a total of 60 oocytes was used (10 oocytes per culture well, three technical and two biological repeats).

##### 2.1.5. Parylene-C coating of BM Flex 80A maintains the fertilization and developmental competence of oocytes

While PC coating of elastomeric materials prevented oocyte degeneration and preserved the functional ability of oocytes to resume and progress through meiosis normally, the ultimate test of egg quality is whether the resulting gamete is capable of supporting fertilization and early preimplantation embryo development. Therefore, to see if PC coating was able to truly maintain egg quality, we performed *in vitro* fertilization (IVF) using eggs matured *in vitro* in BM 80A + PC or in a standard polystyrene 24-well plate (IVM control) and tracked fertilization and preimplantation development (**Figure 5A**). It is well established that eggs matured *in vivo* have superior developmental competence compared to those matured *in vitro*.*^[37,38]^* Hence, to ensure that IVF conditions were optimal, *in vivo* MII oocytes were used; 100% of these eggs developed to the blastocyst stage (**Figure 5A, Figure S3, Supplementary Video S1)**.

**Figure 5.**
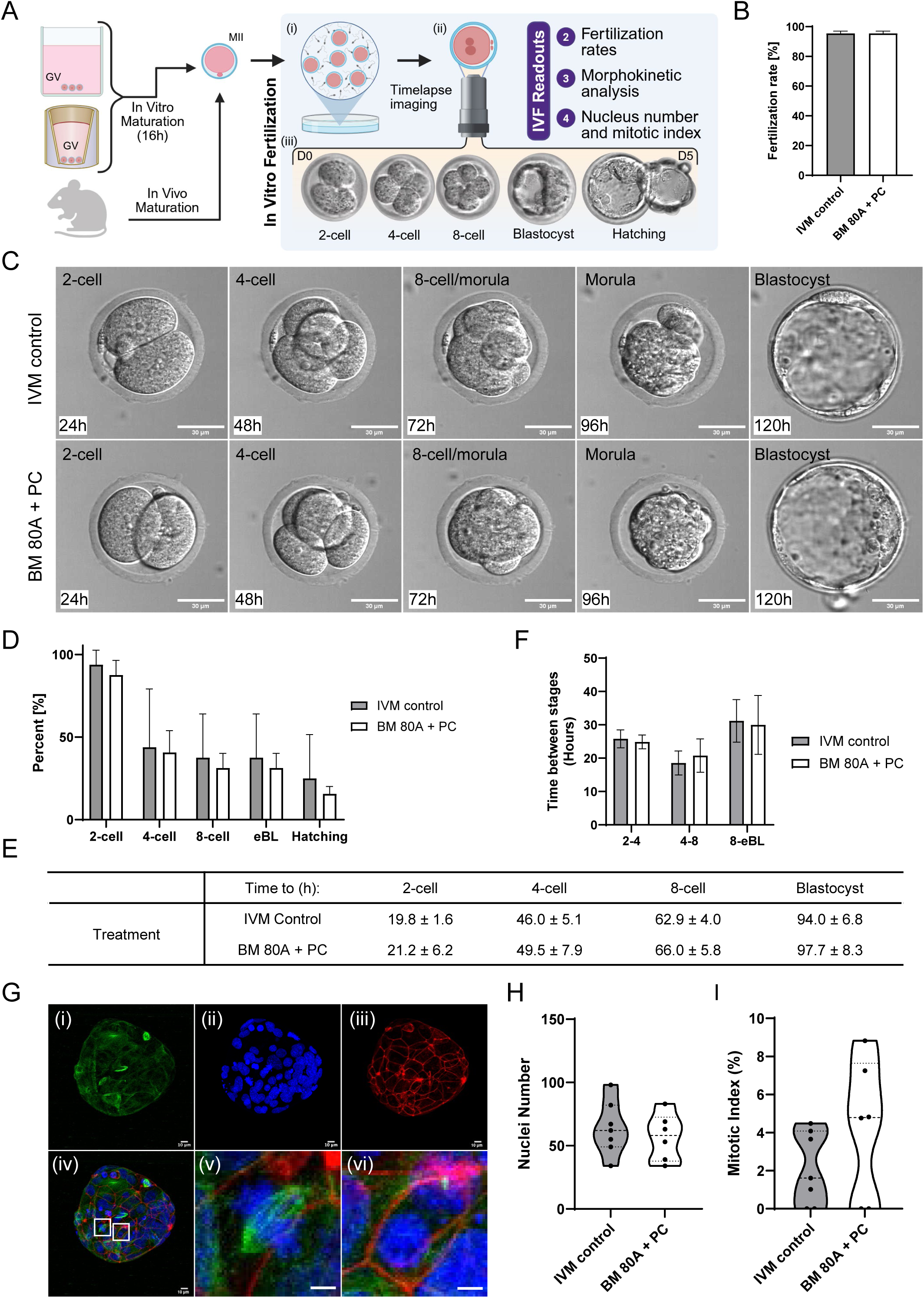
Baseline morphokinetic parameters of embryo development. A) Healthy MII oocytes from either IVM control, PC coated BM 80A or from *in vivo* mice are (i) fertilized during the *in vitro* fertilization (IVF) verification stage. (ii) If a pronucleus or second polar body is formed it is cultured for 5 days and imaged continuously using timelapse imaging. (iii) During embryo culture, the zygote progresses from the 2-cell until blastocyst or hatching stage. IVF readouts utilize metrics for developmental competence, including fertilization rates, morphokinetic analysis and mitotic index. B) Fertilization rates (n=109 and n=112 oocytes for IVM and PC, respectively). C) Representative timelapse images with 24-hour increments for embryos that reached blastocyst stage from IVM control and PC coated BM 80A oocytes (n=2, 16 embryos each). D) Developmental rates for each condition. Quantification of the number of embryos reaching 2-cell, 4-cell, 8-cell and early blastocyst (eBL) stages (n=2, 16 embryos). E) Morphokinetic timepoint analysis of embryo development (n=2, 16 embryos). F) Calculation of time between successive stages up to early blastocyst (n = 12 for control and n = 7 for PC). G) Representative z-projections of day-5 embryos stained with actin (rhodamine phalloidin, red), α-tubulin (green), and DNA (DAPI, blue) by immunocytochemistry. scale bar: 10 µm. H) Quantification of DAPI-positive nuclei counts (n=7 for IVM control and n=6 for BM 80A + PC). I) Mitotic index percentage of day-5 embryos (n=7 for IVM control and n=6 for BM 80A + PC).

Similar to our previous findings, all the oocytes matured in uncoated BM 80A were degenerate, whereas 96.7 *±* 2.4% and 95.8 *±* 3.5% of those matured in control conditions or with BM 80A + PC reached the MII stage, respectively. Fertilization was highly efficient for MII eggs derived from both control or BM 80A + PC conditions, with 95.4 *±* 1.6% and 95.4 *±* 1.5%, respectively reaching the pronuclear stage at 6 hours post-insemination (**Figure 5B**). A total of 32 fertilized eggs per condition were then imaged using timelapse microscopy to monitor development to the 2-cell, 4-cell, 8-cell, and blastocyst stages (**Figure 5C-D, Supplementary Videos S2 and S3)**.

We first compared the ratio of embryos reaching each of these developmental stages, and a high number of embryos reached the 2-cell stage (93.8 ± 8.8% for IVM control vs 87.5 ± 8.8% for BM 80A + PC). This was followed by a reduction of progressing embryos reaching the 4-cell stage (43.8 ± 35.4% vs 40.6 ± 13.3%), a high percentage of those then reached 8-cell (37.5 ± 26.5% vs 31.3 ± 8.8%), early blastocyst (eBL, 37.5 ± 26.5% vs 31.3 ± 8.8%), and finally hatching (25.0 ± 26.5% vs 15.6 ± 4.4%) stages (**Figure 5D-E**). The resulting blastocysts across conditions had normal morphology with clear inner cell masses and expanded blastocoel cavities.

This was followed by a morphokinetic analysis, where we compared the developmental velocity between stages. No significant difference in time intervals between successive cleavage divisions of both groups was seen: 2- to 4-cell (25.8 ± 2.7 h for IVM control vs 24.9 ± 2.1 h for BM 80A + PC), 4- to 8-cell (18.6 ± 3.6 h vs 20.8 ± 5.0 h), and 8-cell to early blastocyst (31.2 ± 6.4 h vs 30.0 ± 8.8 h) (**Figure 5F**).

Lastly, we compared the cell number and mitotic index of the embryos after 5 days of culture (**Figure 5G-I**). Blastocyst cell number is a well-established marker of developmental and implantation potential.*^[39,40]^* The mitotic index, or the percentage of actively dividing cells in the embryo, is a further readout of embryo health and developmental potential.*^[41]^* Blastocysts from IVM control and BM 80A + PC were similar in cell number to each other, counting an average of 63.9 ± 21.2 and 56.8 ± 18.6 (**Figure 5H**). The IVM control mitotic index (2.1 ± 1.9%) did not differ significantly from BM 80A + PC, which was 4.3 ± 3.7% (**Figure 5I**). Together, these findings indicate that Parylene-C coating of BM 80A renders this toxic material completely biocompatible, as the developmental competence of oocytes was not affected compared to IVM control.

### 3. Discussion

The Multi-Endpoint Oocyte Safety Assay (MEIOSA) revealed a range of toxicity profiles caused by four biocompatible SLA-printed resins, varying from delayed meiotic progression to complete oocyte degeneration. Despite their ISO 10993-based classification as biocompatible, none of the resins tested in this study passed the MEIOSA standard after regular post-processing done according to manufacturer protocols. However, we demonstrated that a 5 µm Parylene-C coating rendered inserts that induced various levels of cytotoxicity completely biocompatible with the oocyte. Remarkably, when one of the most toxic materials was coated with Parylene-C, the resulting oocytes exhibited the same fertilization capacity and preimplantation embryo developmental potential as oocytes that were matured in standard conditions.

MEIOSA is a multidimensional biocompatibility screening method that goes beyond standard live/dead evaluation. Current methods to determine if a material is biocompatible rely on viability-based approaches using the ISO 10993 standards, which do not effectively predict if these materials should be used to engineer MPS. For example, HL-1 cardiomyocytes cultured in Clear V4 after sonication in IPA had a cell viability of 92% ± 0.8%.^[36]^ In another study, BM 50A and BM 80A were determined to be biocompatible using ISO 10993-5 based methods because LDH secretion by human dermal fibroblasts was below the polystyrene reference.^[42]^ Viability-based endpoints also fail to detect sublethal perturbations of cell behavior. For example, transcriptomic profiling of adipose-derived stem cells exposed to leachates from BioMed Clear and Clear V4 resins revealed downregulation of extracellular matrix genes and upregulation of cell adhesion and lipid metabolism pathways.^[25]^ Moreover, leachates from biocompatible 3D-printed polymers can act as endocrine disruptors, elevating estrogen receptor (ER) transactivation in a cell-based ER-mediated bioassay.^[24]^ Of note, none of the resins in these examples were biocompatible within MEIOSA, demonstrating its sensitivity and ability to be a rapid phenotypic biocompatibility screening assay that provides insights beyond canonical live/dead metrics. Zebrafish embryo teratogenicity and *in vitro* bovine fertilization assays have also been utilized successfully to evaluate the effects of 3D print leachates.^[20,21,24]^ However, these embryo cultures take up to 8 days to complete, so this method is advantageous as it allows for efficient screening within 16 hours. *In vitro* bovine oocyte maturation and fertilization assays have been used for toxicity screening broadly, as they show similarities in their maturation process with humans and reduce the need for laboratory animal use.^[43–45]^ Our current model relies on mouse oocytes, sperm, and embryos, which have some temporal and morphological differences to the human.^[44]^ Furthermore, although hyperstimulation protocols maximize the oocyte yield per mouse, oocytes are a biolimited resource. An ideal solution would be stem-cell-derived models, and models that focus on different stages and features of the embryo have been developed for DART testing purposes.^[46–48]^

The ability of MEIOSA to distinguish degrees of toxicity becomes apparent when comparing rigid and elastomeric ISO 10993-certified resins. Rigid BioMed resins (BM Clear and BM Durable) did not impact oocyte survival but severely compromised meiotic progression and resulted in abnormal phenotypes. Overcuring protocols improved the biocompatibility of BM Clear, but efficiency will ultimately depend on factors including print dimensions and length of culture. Elastomeric BioMed resins (BM Elastic 50A and BM Flex 80A) were acutely cytotoxic, causing 100% oocyte degeneration early in the 16-hour culture period. Many leachates from 3D-printed resins could be responsible for the oocyte-based toxicity. For example, a previous study determined that Tinuvin 292 light stabilizer in Dental LT was ovotoxic.^[11]^ The stark increase in cytotoxicity of BM 50A and BM 80A, relative to the more rigid biomaterials, is likely due to plasticizers which are added to maintain elastomeric properties.^[49]^ Interestingly, phthalates are a common category of plasticizers that have demonstrated reproductive toxicity and have been found in leachates of SLA printable resins.^[24,28,50]^ However, these commercial resins have proprietary compositions, so their exact chemical makeup remains unknown. Identification of the leachates originating from these resins is an important area of future investigation.

While there are multiple ways to reduce the toxicity of 3D printed polymers, Parylene-C has proven to be an effective barrier to revert the effect of even the most cytotoxic resins, which was validated by IVF. Common methods to improve biocompatibility include the development of custom resin formulations that utilize less toxic components,^[51–53]^ application of heat,^[25,26]^ use of plasma cleaning,^[11]^ optimization of curing conditions to improve crosslinking,^[20,36]^ removal of residual compounds via washing,^[14,35,54–57]^ use of coatings that delay leaching,^[21]^ or a combination of these strategies.^[36,58,59]^ Parylene-C was evaluated as a possible universal solution to prevent any leachates from entering the culture medium. It is a common and affordable FDA USP Class VI polymer able to form deformable and conformal coatings that uniformly cover intricate 3D-printed geometries at room temperature without the use of solvents, catalysts, or plasticizers.^[33,60,61]^ Parylene-C also presents long-term thermal stability as no negative impact was observed when used in neural sensors implanted *in vivo* for up to one year.^[62,63]^ Furthermore, iPSC-derived neural cells have been cultured for 14 days with PC coated resin printed inserts without negative effect on viability.^[64]^ Importantly, Parylene-C effectively coats microfluidic channels in a linear microfluidic model and prevents cytotoxicity in sensitive primary cells.^[33]^ Parylene-C coating has also been shown to minimize adsorption of lipophilic compounds and preserve cell viability and phenotypic fidelity in a PDMS-based feto-maternal interface MPS utilizing human decidual and amnion epithelial cells, confirming its compatibility with reproductive tract somatic cells. ^[61]^

Biocompatibility of MPS with reproductive cells is particularly important, as reproductive MPS could bridge the significant anatomic, developmental, and endocrine differences between common animal models and humans.^[65–67]^ Oocytes and embryos have been used prior to test the biocompatibility of 3D printed materials, but this is the first study to our knowledge that utilizes both gamete and zygote to test resins and the effectiveness of Parylene-C coating.^[11,20,21,68]^ In this study, cytotoxic compounds were only allowed to leach out over a 40-hour period (24 hours incubation and 16 hours of IVM), and future experiments may need to determine the relation between PC thickness and maximal culture period. The PC barrier function seems long-term, as leaching experiments have shown that leachate levels drop 50-fold with a 10 µm PC coating after 70 days of incubation.^[34]^ Nonetheless, a 5 µm Parylene-C coating showed excellent biocompatibility properties, rescuing meiotic competence across all materials. To test if oocytes were able to maintain fertilization and preimplantation developmental potential, IVF was used. Here, oocytes cultured with PC-coated BM Flex 80A, a resin that normally induces immediate cell death, showed no difference in fertilization efficiency, embryo growth or cell number and mitotic index after 5 days of culture. It is possible that genetic or epigenetic testing of the day 5 embryo will give more mechanistic insights than the mitotic index, which can be evaluated in future studies. Alternatively, further insights from morphokinetic analysis into specific pathways can be accelerated using convolutional neural networks and artificial intelligence.^[69]^

These advances enable the use of previously toxic resin-3D-printed materials for MPS containing sensitive cultures and germ cells, not only advancing efforts toward a reproductive-tract-on-a-chip but also expanding the toolbox of materials available for MPS more broadly.^[66,67]^

### 4. Conclusion

We validated MEIOSA as a highly sensitive, multi-endpoint screening assay that generates concrete results within 16 hours and can be followed up with IVF-based validation. Using this approach, we demonstrated that four ISO 10993-certified “biocompatible” BioMed resins induce cytotoxic effects to the oocytes which could be rescued using a 5 µm-thick coating of Parylene-C. These findings validate MEIOSA as a high stringency biocompatibility standard for 3D-printed microphysiological systems and the use of conformal Parylene-C coating to decouple material selection from biocompatibility constraints.

### 5. Experimental Section/Methods

#### Animals

Reproductively young adult CD-1 female mice were obtained from Envigo (Indianapolis, IN, USA). Mice were kept in barrier facilities under temperature, humidity, and light control (14 h light/10 h dark cycles) with food and water provided ad libitum. Mice were allowed to acclimatize for at least a week before being used for experiments. Water and Teklad Global irradiated 2916 chow containing minimal phytoestrogens and no soybean or alfalfa meal were provided ad libitum. All experimental protocols and animal procedures were approved by the Institutional Animal Care and Use Committee (IACUC) of Northwestern University and were performed in accordance with National Institutes of Health Guidelines.

#### Three-dimensional design and post-processing

Culture inserts were designed using Autodesk Fusion360 (v.2.0.21508, Autodesk Inc., CA, USA) and imported into PreForm 3D-print preparation software (Formlabs Inc, MA, USA). Models were printed with five different resins (Clear V4, BioMed Clear, BioMed Durable, BioMed Elastic 50A, and BioMed Flex 80A) using a Form 3B+ SLA printer at the thinnest layer thickness available (Formlabs Inc, MA, USA, Figure 1C). Printed inserts were thoroughly sprayed with 99% isopropyl alcohol, washed and cured using a FormWash and FormCure (Formlabs Inc, MA, USA), respectively, according to manufacturer guidelines (Figure 1C).

#### Parylene-C (PC) coating and sterilization

PC coatings were applied using an SCS Labcoater Parylene Deposition System (Specialty Coating Systems Inc, IN, USA). Prior to deposition, inserts were cleaned via sonication in 99% isopropanol (IPA) for 5 minutes and then transferred into a fresh 99% IPA bath for approximately 10 minutes until loading. A total of 21.05g of parylene-C dimer was used, producing an estimated coating thickness of 5 µm. During deposition, the furnace temperature was raised to 690 °C and the vaporizer to 180 °C. Deposition initiated around 135–140 °C, with a chamber pressure of approximately 35 Torr maintained during active coating. Once the PC was exhausted, the system automatically transitioned into a pump-down phase, returning to 1 Torr. The deposition process lasted approximately 3.5 hours at a substrate temperature of 55 °C. The width of coated and uncoated inserts was measured using a caliper. All inserts, whether PC coated or uncoated, were sterilized in the biosafety hood via submersion in 70% isopropanol (IPA) for 5 minutes and were then airdried for 30 minutes with UV light exposure.

#### Overcuring protocols (OCP)

BM Clear inserts were processed after printing according to the manufacturer’s standard protocol (Figure 1C). Following this, inserts were (i) left untreated, (ii) received additional UV curing in Milli-Q water at 70 °C for 2 hours (OCP1), or (iii) UV curing in Milli-Q water at 70 °C for 1 hour followed by incubation in a 70 °C oven for 16 hours (OCP2). Inserts were subsequently sterilized and prepared for downstream *in vitro* maturation (IVM) culture experiments.

#### Isolation and in vitro maturation (IVM) of oocytes

Germinal vesicle (GV) oocytes were collected from 6–12 weeks old CD-1 mice using previously published methods.^[70]^ In summary, Female CD-1 mice were hyperstimulated with 5 IU pregnant mare serum gonadotropin (PMSG, ProSpec-Tany TechnoGene, NJ, USA, Cat No. HOR-272) injection to maximize oocyte yield. Ovaries were harvested 44-46 hours post-injection and cumulus-oocyte-complexes (COCs) were isolated with insulin syringes, cumulus cells were then mechanically removed with a 100 µm stripper tip to collect denuded oocytes. Prior to IVM culture, 500 µL of culture media consisting of α-MEM Glutamax supplemented with 3 mg/mL bovine serum albumin (BSA) and 0.5% penicillin-streptomycin (PS, Life Technologies) was conditioned for 24 h at 37 °C in a humidified atmosphere of 5% CO_2_ in a polystyrene 24-well with or without inserts (Uncoated and PC coated Clear V4, BM Clear, BM Durable, BM 50A, and BM 80A). For IVM followed by IVF, 10mg/ml of fetal bovine fetuin (Sigma, F-3385) is added to the culture media to prevent zona pellucida hardening of oocytes during IVM. During IVM, 10 oocytes were seeded in each well (n=3 per condition) and were cultured for 16 hours. A total of 1,149 oocytes were used to test 11 conditions: a control group (no insert) and five resins, each evaluated in the presence and absence of a PC coating.

#### Meiotic progression analysis

Oocytes were imaged before and after IVM at 10X, and 20X magnification with a EVOS M7000 imaging system (Thermo Fisher Scientific, US). After IVM, oocytes were evaluated for their meiotic stage based on established morphological criteria.^[11]^ Oocytes with a visible, intact germinal vesicle (GV) were classified as arrested in prophase of meiosis I (GV). Those that had lost the GV but had not yet extruded a polar body were considered to have undergone germinal vesicle breakdown (GVBD) and were in metaphase I (MI). Oocytes that lacked a GV and had extruded a polar body were classified as arrested at metaphase II (MII). Oocytes were categorized as abnormal if internal aggregates, membrane protrusions and large vacuole-like bubbles were present. Oocytes that appeared shrunken, flat, or darkened were classified as degenerate.

#### Chromosomal alignment analysis

After IVM, oocytes that were not degenerate were fixed in 3.8% paraformaldehyde (Electron Microscopy Sciences, Hatfield, PA, USA) containing 0.1% Triton X-100 (Sigma-Aldrich) for 25 minutes at 37 °C. Following fixation, they were washed four times for 5 minutes each in a blocking buffer composed of 1× PBS, 0.01% Tween-20, and 0.3% BSA (all from Sigma-Aldrich). The oocytes were then incubated in a permeabilization solution (1× PBS, 0.1% Triton X-100, and 0.3% BSA) for 15 minutes at room temperature, followed by two additional 5-minute washes in blocking buffer. To analyze spindle morphology, immunofluorescence was performed with an anti-tubulin antibody as described previously.^[71]^

Confocal imaging was performed on a Leica SP5 inverted laser scanning confocal microscope (Leica Microsystems, Wetzlar, Germany) at 40X objective, using 405, 488, and 543 nm laser lines. Images were processed with LAS AF software (Leica Microsystems) and analyzed using FIJI (NIH, Bethesda, MD, USA). Spindles oriented perpendicular to the image plane were excluded. Chromosome alignment at the metaphase II was scored as 0 (fully aligned chromosomes), 1 unaligned chromosome, >1 unaligned chromosome.

#### In vitro fertilization

Preparations for *in vitro* fertilization (IVF) were made the day before: EmbryoMax® HTF was supplemented with BSA to 4 mg/mL (HTF+BSA) for sperm capacitation and insemination. To obtain mature eggs for IVF (*in vivo* control), female mice were super ovulated with an intraperitoneal (i.p.) injection of 5 IU PMSG, followed by an i.p. injection of 5 IU human chorionic gonadotropin (hCG) 44–46 hours later. COCs were collected from the oviduct 14–16 hours post-hCG injection and loaded per fertilization well. Insemination (IVM control, BM 80A + PC, and *in vivo* control) was performed with capacitated sperm to a final concentration of 1 × 10^6^ cells/ml for 6 hours. Fertilized eggs were identified by two pronuclei (2PN) and were loaded in the EmbryoSlides (Vitrolife, Denver, CO) containing KSOM (EMD millipore) medium. EmbryoSlide with culture media was overlaid with 1.6 ml of mineral oil (Sigma-Aldrich, St. Louis, MO) and equilibrated in the EmbryoScope+™ for 9–24 h. To assess the morphokinetic, we assessed the morphological parameters of 2-cell, 4-cell, 8-cell, and blastocyst stages manually on EmbryoViewer.

#### Blastocyst cell counts

After 5 days of embryo culture, all blastocysts were fixed in 3.8% paraformaldehyde for 1 hour at room temperature and rinsed twice in blocking buffer (PBS with 0.3% BSA, and 0.01% Tween-20). Then, permeabilized for 15 minutes in phosphate-buffered saline (PBS) supplemented with 0.3% bovine serum albumin (BSA), 0.1% Triton X-100, and subsequently rinsed twice in blocking buffer. To preserve the integrity of the blastocoel cavity, blastocysts were sequentially transferred through a graded series of Vectashield solutions containing DAPI (25%, 50%, 75%, and 100%; Vector Laboratories, Burlingame, CA, USA). Imaging was performed as described in chromosomal alignment analysis. Image analysis was conducted with FIJI, and cell counts were determined based on the number of DAPI-stained nuclei. Optical sections for each embryo were analyzed using ImageJ, and the number of cells per blastocyst was calculated based on the number of DAPI-positive nuclei. Each nucleus was tracked through optical sections to avoid double counting. The mitotic index was calculated as the number of blastomeres in metaphase or anaphase (as evidenced by the DNA configuration) divided by the total number of cells in the embryo.

#### Statistical Analysis

Data are presented as the mean ± standard deviation, and each experiment was repeated twice using a minimum of 30 oocytes per experiment. Unless otherwise noted, error bars represent the standard deviation. All statistical analyses were performed using GraphPad Prism 10.0 (GraphPad Software Inc., San Diego, CA, USA). The normal distribution of data was evaluated with the Shapiro–Wilk test and outliers were identified using the ROUT method. Analysis between groups was performed with Student’s *t*-test or Mann–Whitney *U* test. Multiple comparisons were analyzed with two-way analysis of variance (ANOVA) test, followed by Tukey’s multiple comparison tests. *P* values <0.05 were considered statistically significant.

## Acknowledgements

This work made use of the EPIC facility (RRID: SCR_026361) of Northwestern University’s NUANCE Center, which has received support from the IIN and Northwestern’s MRSEC program (NSF DMR-2308691). Figures were created with BioRender.com.

## Supporting Information

**Figure S1.**
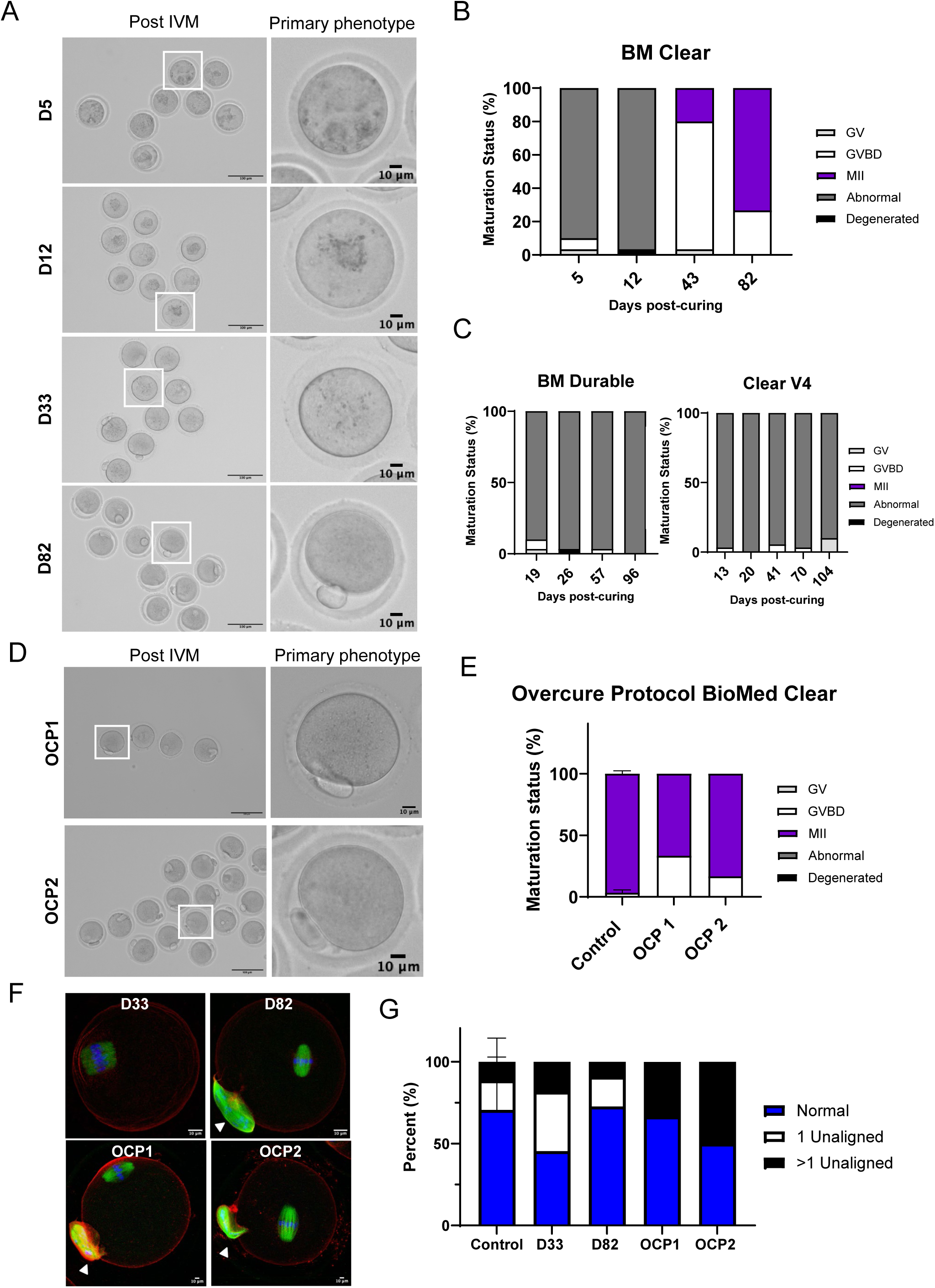
Effect of aging and overcuring protocols on BioMed Clear. A) Representative images of oocytes observed after IVM (left, scale bar: 100 µm) with primary phenotypes (right, scale bar: 10 µm) for BM Clear at different timepoints post-curing. B) Resulting meiotic progression incidence for BM Clear at different timepoints post-curing. For each timepoint 30 oocytes were used. C) Meiotic progression incidence for BM Durable and Clear V4 at different timepoints post-curing. For each timepoint 30 oocytes were used. D) Representative images of primary phenotype observed after IVM for BM Clear after overcure protocol 1 and 2 (OCP). OCP 1 and OCP 2 conditions were tested using 15 and 60 oocytes, respectively. E) Resulting meiotic progression incidence for BM Clear after OCP 1 and 2 compared to control. F) Representative z-projections of MII eggs. Actin (rhodamine phalloidin, red), α-tubulin (green), and DNA (DAPI, blue) were detected by immunocytochemistry. PB indicated by triangle, scale bar: 10 µm. G) Incidence of normal and aberrant chromosome alignment for aged and OCP 1 and 2 treated BM Clear.

**Figure S2.**
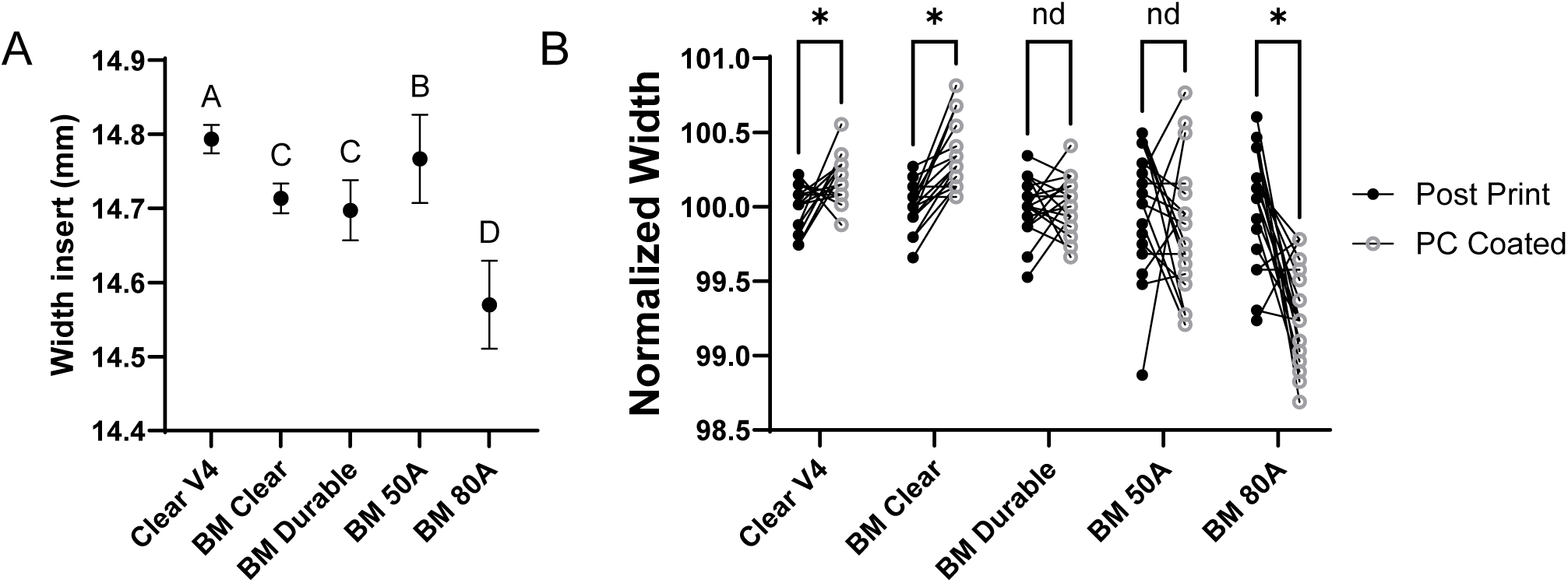
Effect of Parylene-C coating procedure on insert dimensions. A) Width measurements of fully cured inserts for each resin type. Compact letter display (A, B, C, D) shows a significance of 0.05 or less between resins (n=54 for Clear v4, BM Clear, BM Durable and n=56 for BM 50A and BM 80A). B) Normalized width after curing and after coating shows differential effect of the coating procedure. * represents *P* < 0.05.

**Figure S3.**
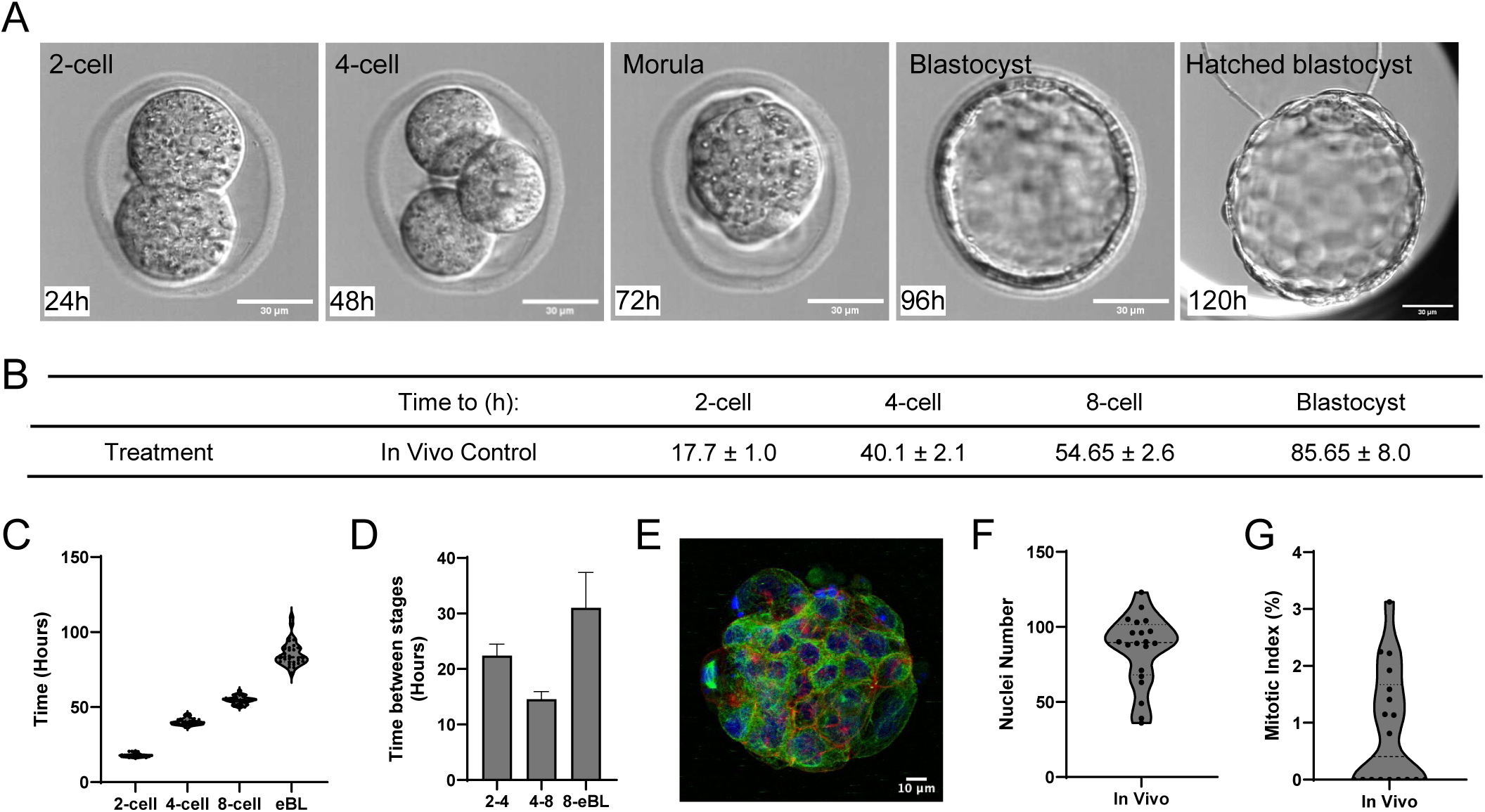
Developmental parameters of embryo derived from *in vivo* obtained MII oocyte. A) Representative timelapse images with 24-hour increments for embryos that reached blastocyst stage. B) Morphokinetic timepoint analysis of embryo development (n=2, 16 embryos). C) Quantification of time required to reach 2-cell, 4-cell, 8-cell, and early blastocyst stages (eBL) of development of healthy MII oocytes from *in vivo* mice (n=2, 16 embryos). D) Calculation of time between successive stages. E) Representative z-projections of day-5 embryo stained with actin (rhodamine phalloidin, red), α-tubulin (green), and DNA (DAPI, blue) by immunocytochemistry. F) Quantification of DAPI-positive nuclei counts (n=20). G) Mitotic index percentage of day-5 embryos (n=18).

### Table of contents entry

New approach methodologies such as microphysiological systems (MPS) are designed to reduce reliance on animal models for basic and clinical research. Leachates from resin-3D-printed components represent a risk to incorporating 3D-printed components into MPS. MEIOSA is a Multi-Endpoint Oocyte Safety Assay that stratifies biocompatibility of ISO 10993 compliant resins and demonstrates the ability of Parylene-C to rescue cytotoxicity.

**Supplementary Video S1.** Representative timelapse video of embryo derived from *in vivo* obtained MII oocyte.

**Supplementary Video S2.** Representative timelapse video of embryo derived from MII oocyte matured *in vitro*.

**Supplementary Video S3.** Representative timelapse video of embryo derived from MII oocyte matured *in vitro* with Parylene-C coated BioMed Flex 80A.

